# Human Messenger RNA Harbors Widespread Noncoding Splice Isoforms

**DOI:** 10.1101/2025.05.06.652571

**Authors:** Qili Shi, Haochen Li, Junjiao Song, Hao Wu, Yu Zeng, Jiawen Jiang, Shengli Li, Zhiao Chen, Xianghuo He

## Abstract

Pre-messenger RNA transcribed from canonical protein-coding genes is frequently alternatively spliced into diverse mature mRNA isoforms, thus translating into various protein products. However, it remains unknown whether human mRNA from a protein-coding gene harbors noncoding splice isoforms in the genome. Herein, we discovered 15,836 mRNA noncoding splice isoforms across 7,298 protein-coding genes in human. mRNA noncoding isoforms are mainly produced by alternative splicing and polyadenylation, which display tissue-specific distributions and involve in RNA processing pathways. Notably, mRNA noncoding isoforms are frequently upregulated in cancer. Differentially expressed mRNA noncoding isoforms are associated with the cancer hallmarks and can independently predict patient survival. These findings unraveled human mRNA harbors widespread noncoding splice isoforms, and highlight the dual characters of mRNA including protein-translation isoforms and noncoding splice isoforms, providing a new dimension to our understanding of mRNA functional property.

## Introduction

The human genome encodes approximately 20,000 protein-coding genes constituting 2%-5% of the total transcriptome. mRNA, transcribed from protein-coding genes, is exported from the nucleus to the cytoplasmic translation machinery, exerts the biological functions through the proteins they encode (*1*). In contrast, the majority (75-90%) of transcriptional output from human genome comprises non-coding RNAs (ncRNAs), which have emerged as sophisticated regulators in the cell (*2*). Recently, a few traditionally considered untranslatable ncRNAs have been shown to possess functional open reading frames (ORFs) capable of producing biologically active peptides (*3*). Notably, several protein-coding mRNAs have been demonstrated as bi-functional molecules. The same individual mRNA molecule exhibited both protein-coding and non-coding functions via the structural domains within 5′ UTR (untranslated region), CDS (coding sequence), and 3′ UTR regions (*4–6*).

Pre-messenger RNA, transcribed from the protein-coding genes, commonly undergoes alternative splicing to generate multiple mRNA isoforms with related, distinct, or even opposing functions. Notably, disease-associated aberrant splicing can redirect pre-messenger RNA processing to produce non-coding splice variants, particularly in cancer (*7, 8*). For instance, Simonelli et al. identified an exon2-lacking variant of DNA repair polymerase β that does not produce a functional protein but instead generates ncRNAs in gastric cancer (*9*). Similarly, the PPP1R10 gene, which encodes the Protein Phosphatase-1 (PP-1) Nuclear Targeting Subunit (PNUTS), expressed lncRNA-PNUTS due to an alternative 3′ splice site in exon 12 that disrupts the PNUTS ORF in breast cancer (*10*). Programmed death-ligand 1 (PD-L1) gene produced a long non-coding RNA isoform PD-L1-lnc by alternative splicing in lung adenocarcinoma (*11*). Beyond human cancer, an intronic promoter within intron 2 in the murine proteinase 3 gene derived the expression of an alternative mRNA transcript that seems not to be translated in mouse leukemia cell (*12*). In addition, UV irradiation could induce functional noncoding splice variants through alternative pre-mRNA processing or polyadenylation (*13, 14*). These advancements demonstrated that aberrant splicing as a molecular switch could convert a protein-coding mRNA isoform into a regulatory ncRNA variant in the disease or stress conditions.

Alternative splicing of protein-coding genes in human genome generates an estimated 110,000 splice isoforms (GENCODE Release 47), which greatly expands proteomic complexity by producing functionally distinct protein isoforms with tissue-specific expression patterns and divergent physiological roles. A previous report analyzed the top-ranked expressed transcripts of human protein-coding genes and mentioned that some transcripts like processed_transcripts or retained_introns of protein-coding genes might belong to noncoding biotypes (*15*). To date, it still remains unknown whether human mRNA splice isoforms from the same protein-coding gene harbor noncoding splice isoforms besides protein-coding counterparts in the genome.

In this study, we systematically identified 15,836 mRNA noncoding splice isoforms across 7,298 protein-coding genes accounting for 36% of total 20, 270 protein-coding genes in human genome, with 64.3% representing previously unannotated isoforms. These mRNA noncoding splice isoforms display higher expression levels than canonical lncRNAs but lower than protein-coding mRNA isoforms. mRNA noncoding splice isoforms are produced by alternative splicing and polyadenylation, which display tissue-specific distributions and are highly related to RNA processing pathways. In cancer, we detected 12,156 mRNA noncoding splice isoforms across 33 tumor types. Differentially expressed noncoding splice isoforms (DEIs) are associated with patient prognosis, and correlated with the established cancer hallmarks. Together, these findings unraveled the dual characters of mRNA transcribed from the same human protein-coding gene: the canonical splice isoforms preserve their protein-translational activities; its noncoding splice isoforms diverge to act as regulatory ncRNAs.

## Results

### Genome-wide characterization of noncoding splice isoforms in human mRNA

To comprehensively assess the coding potential of full-length RNA transcripts within human protein-coding gene loci, we first retrieved 403,693 transcript isoforms expressed in human normal tissues from the FLIBase database, generated using third-generation long-read sequencing technologies (*16*). To improve reliability and tissue relevance, we filtered these isoforms based on RNA-seq expression data from the GTEx project, retaining transcripts with a median TPM > 0 in at least one tissue. Genomic coordinates were assigned using the Ensembl GRCh38 annotation, and only transcripts located within protein-coding gene loci were included. Coding potential was evaluated using three independent tools: CPAT (Coding Potential Assessment Tool) (*17*), CPC2 (Coding Potential Calculator 2) (*18*), and GeneMark (*19*). Transcripts consistently predicted to be non-coding by at least two of the three tools were retained, resulting in a total of 22,318 noncoding transcripts. These noncoding transcripts were further classified into two categories: 6,482 were annotated as canonical noncoding RNAs, including lncRNAs, antisense RNAs, and processed transcripts overlapped with protein-coding gene loci, while the remaining 15,836 noncoding transcripts were annotated as mRNA noncoding splice isoforms, which are produced by protein-coding gene, contain multiple exons and share at least one identical exon with the annotated protein-coding mRNA isoforms from the same protein-coding gene. Of these, 10,187 (64.3%) were newly identified and annotated (**Fig. 1A, table S1**).

**Fig 1.**
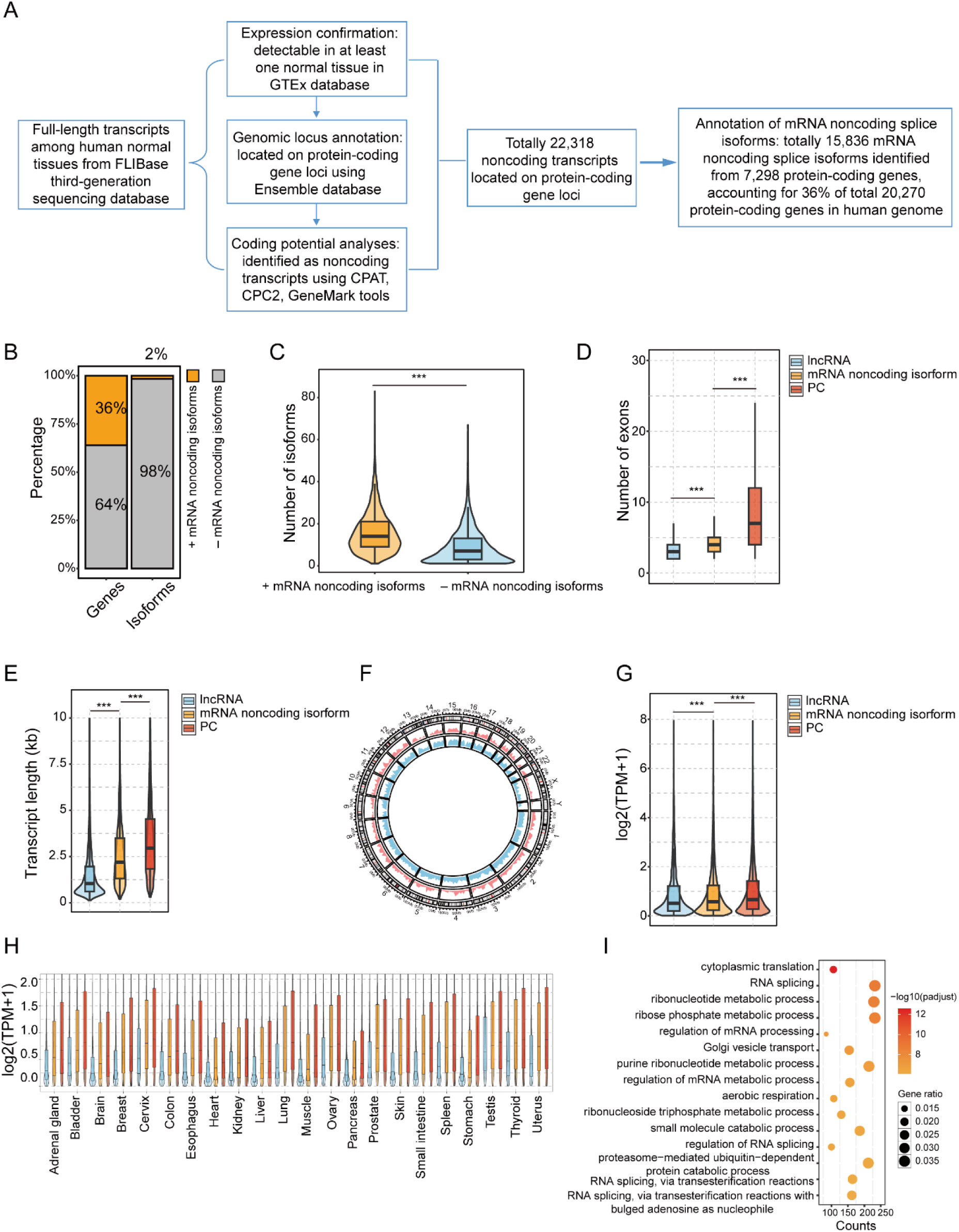
Genome-wide characterization of noncoding splice isoforms in human mRNA. **A.** Schematic overview of the workflow used to identify and classify mRNA-derived noncoding isoforms, including transcriptome annotation filtering, coding potential evaluation, and transcript categorization. **B.** Stacked bar plots showing (left) the proportion of protein-coding genes that generate at least one mRNA noncoding isoform, and (right) the relative proportions of mRNA noncoding isoforms among all transcript isoforms. **C.** Box and violin plots comparing the total number of transcript isoforms per gene between protein-coding genes with and without mRNA noncoding isoforms. Statistical significance was calculated using the Wilcoxon test. **D.** Box and violin plots showing the distribution of exon counts among mRNA noncoding isoforms, canonical lncRNAs, and protein-coding isoforms (PC). Statistical significance was calculated using the Wilcoxon test. **E.** Box and violin plots displaying transcript length distributions of mRNA noncoding isoforms, canonical lncRNAs, and protein-coding isoforms (PC). Statistical significance was calculated using the Wilcoxon test. **F.** Circular plot showing the genomic distribution of mRNA noncoding isoforms and protein-coding genes across human chromosomes. The outer ring displays chromosome ideograms; inner tracks indicate gene (blue) and transcript (red) densities. **G.** Box and violin plots showing expression levels (TPM) of mRNA noncoding isoforms, canonical lncRNAs and protein-coding isoforms (PC) across human normal tissues. Statistical significance was calculated using the Wilcoxon test. **H.** Box plots summarizing average expression levels (TPM) of mRNA noncoding isoforms, canonical lncRNAs and protein-coding isoforms across multiple tissues. **I.** Scatter plot presenting functional enrichment results for protein-coding genes producing noncoding isoforms. Each dot represents an enriched Gene Ontology term, with dot size indicating gene ratio and color denoting statistical significance. ***, p < 0.001.

These mRNA noncoding splice isoforms originate from 7,298 protein-coding genes, representing approximately 36% of all protein-coding genes (**Fig. 1B**). Genes harboring noncoding splice isoforms exhibited significantly greater isoform diversity than those without, suggesting a strong association between alternative splicing complexity and the presence of noncoding isoforms (**Fig. 1C**). Structural analysis revealed that these noncoding splice isoforms had intermediate exon numbers and transcript lengths compared to canonical lncRNAs and protein-coding transcript isoforms (**Fig. 1D and 1E**). Chromosomal mapping showed that mRNA noncoding isoforms broadly distribute throughout the genome without evident chromosomal bias, and mirror the genomic distribution of protein-coding genes (**Fig. 1F**). Expression profiling across normal tissues revealed that mRNA noncoding isoforms were more abundantly expressed than canonical lncRNAs, but less than protein-coding mRNA isoforms, with similar expression trends across different tissues (**Fig. 1G and 1H**). Functional analyses of the 7,298 protein-coding genes producing noncoding splice isoforms showed significantly enriched in RNA processing pathways, suggesting that these noncoding splice isoforms may exert essential molecular functions related to RNA regulation (**Fig. 1I**).

### Alternative splicing and alternative polyadenylation events in mRNA noncoding isoforms across human normal tissues

Next, we systematically analyzed post-transcriptional RNA splicing events, particularly alternative splicing (AS) and alternative polyadenylation (APA) in mRNA noncoding isoforms originated from protein-coding genes across human normal tissues. Seven distinct AS event types were identified: exon skipping (SE), mutually exclusive exons (MX), retained introns (RI), alternative 3’ splice sites (A3), alternative 5’ splice sites (A5), alternative first exons (AF), and alternative last exons (AL) (**Fig. 2A**). On average, we detected approximately 5,286 AS events involving mRNA noncoding isoforms per tissue, suggesting that AS extensively contributes to generating these noncoding transcripts. Among these, AF events were the most frequent, followed by RI and AL events. We categorized noncoding isoforms based on their AS inclusion ratios into three groups: low (< 0.4), medium (0.4 ≤ inclusion ratio < 0.6), and high (≥ 0.6) (**Fig. 2B**). Approximately 30% of mRNA noncoding isoforms fell into the high inclusion group, indicating frequent involvement of alternatively spliced exons or introns (**Fig. 2C**). Most mRNA noncoding isoforms were associated with a single type of AS event across the 22 tissues analyzed (**Fig. 2D**). Notably, some mRNA noncoding isoforms exhibited tissue-specific AS patterns. For example, transcript *TCONS_00982193* contained an additional exon compared to transcript *TCONS_00982202*, representing a liver-specific SE event (**Fig. 2E**). In parallel, APA events were examined for each mRNA noncoding isoform. Noncoding splice isoforms generally displayed shorter polyadenylated tails compared to other transcripts from the same or different protein-coding genes (**Fig. 2F**). In tissues such as liver, stomach, and muscle, a substantial proportion of noncoding isoforms showed shortened polyadenylation sites. Conversely, in gender-associated tissues such as testis, cervix uteri, and uterus, noncoding isoforms predominantly exhibited lengthened polyadenylation tails (**Fig. 2G**). Liver-specific lengthening of polyadenylation tails was also observed for several noncoding isoforms (**Fig. 2H**). For instance, noncoding isoform *TCONS_00276302* exhibited an extended polyadenylation tail specifically in the liver, correlating with its liver-specific expression (**Fig. 2I**). Together, these findings highlight the extensive impact of AS and APA mechanisms on the generation and regulation of mRNA noncoding splice isoforms across diverse human normal tissues.

**Fig. 2.**
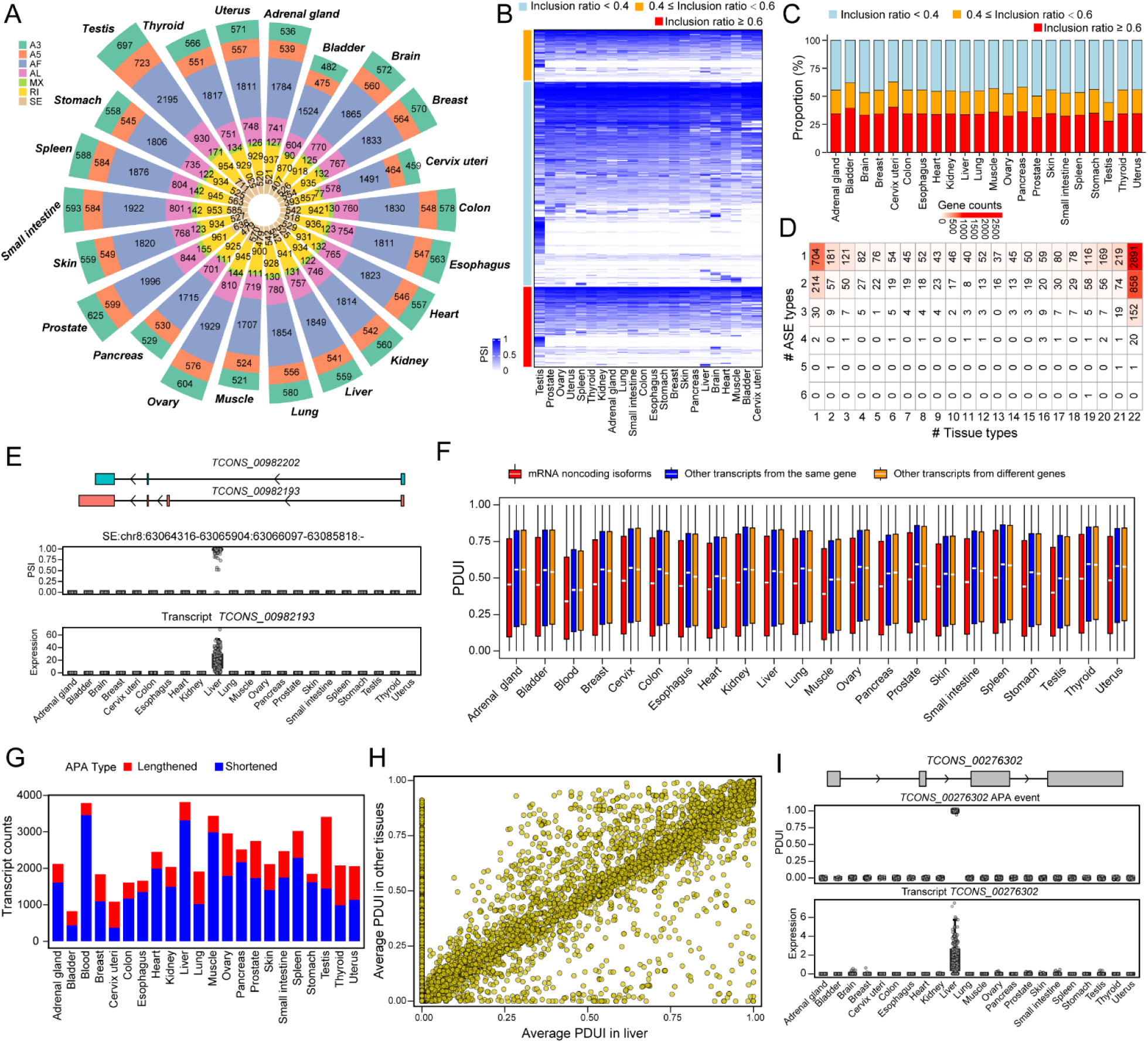
Alternative splicing and alternative polyadenylation events in mRNA noncoding isoforms across human normal tissues. **A.** Circular stacked bar plots depicting the counts of various alternative splicing (AS) event types in mRNA noncoding isoforms across different human normal tissues. **B.** Heatmap illustrating the expression patterns of mRNA noncoding isoforms, classified into high (inclusion ratio ≥ 0.6), medium (0.4 ≤ inclusion ratio < 0.6), and low (inclusion ratio < 0.4) inclusion groups across different human normal tissues. **C.** Stacked bar plots summarizing the proportions of the noncoding isoform inclusion categories shown in panel B across different human normal tissues. **D.** Heatmap revealing the distribution of the number of distinct AS event types observed across varying tissue numbers. **E.** Example of a liver-specific AS event: top, genomic structures of noncoding isoforms *TCONS_00982202* and *TCONS_00982193*; middle, percent spliced-in (PSI) values across different human normal tissues; bottom, expression profile of noncoding isoform *TCONS_00982193*. **F.** Box plots comparing proximal-to-distal polyadenylation usage index (PDUI) values of mRNA noncoding isoforms from protein-coding genes to those from the other transcripts within the same genes and from different genes across different human normal tissues. **G.** Stacked bar plots indicating the number of noncoding isoforms shortened or lengthened by APA events in each tissue. **H.** Scatter plot comparing the average PDUI values between liver and other tissues for each APA event. **I.** Example of a liver-specific APA event: top, genomic structure of transcript *TCONS_00276302*; middle, PDUI values of the APA event across different human normal tissues; bottom, expression profile of noncoding isoform *TCONS_00276302*.

### Tissue-specific expression and functional characteristics of mRNA noncoding splice isoforms across human normal tissues

Furthermore, we assessed the expression variability and specificity of mRNA noncoding isoforms across different human normal tissues. mRNA noncoding isoforms demonstrated considerable expression diversity, classifiable into tissue-ubiquitous, intermediately specific, and tissue-specific expression categories (**Fig. 3A**). To quantify tissue-specific expression, a specificity score was calculated for each noncoding isoform based on comparative expression analysis. mRNA noncoding isoforms exhibited specificity scores similar to those of protein-coding transcripts and other transcripts from the same genes, but lower than those of transcripts originating from noncoding genes (**Fig. 3B**). Some tissue-specific noncoding isoforms originated from tissue-specific genes, either within the same tissues or in different tissues (**Fig. 3C**). For instance, both gene *SPP2* and its associated noncoding isoform *TCONS_00598919* showed liver-specific expression patterns (**Fig. 3D**). In contrast, the gene *FABP6* exhibited small intestine-specific expression, while its noncoding splice isoform *TCONS_00829023* was specifically expressed in the brain (**Fig. 3E**). Moreover, we explored whether the genes expressing these mRNA noncoding isoforms were linked to human embryonic development. Our analysis identified 80 noncoding isoforms associated with embryonic development-related genes: 32 were ubiquitously expressed, 31 intermediately specific, and 17 tissue-specific (**Fig. 3F**). To elucidate potential functional implications of these noncoding isoforms, we conducted correlation analyses between noncoding isoform expression and protein-coding gene expression. Positively correlated protein-coding genes were significantly enriched in RNA processing pathways, including “mRNA processing”, “RNA splicing”, and “ncRNA processing”. Conversely, negatively correlated genes primarily participated in ribosome-associated processes, i.e. “ribonucleoprotein complex biogenesis” and “ribosome biogenesis” (**Fig. 3G**). Additionally, we revealed RNA-binding proteins (RBPs) significantly correlated with noncoding isoform expression, particularly the RBPs such as SRRT, SUGP2, and SMG9 exhibited positive correlations (**Fig. 3H**). Overall, these findings highlight distinct tissue-specific expression patterns and possible functional roles of mRNA noncoding isoforms from protein-coding genes across human normal tissues.

**Fig. 3.**
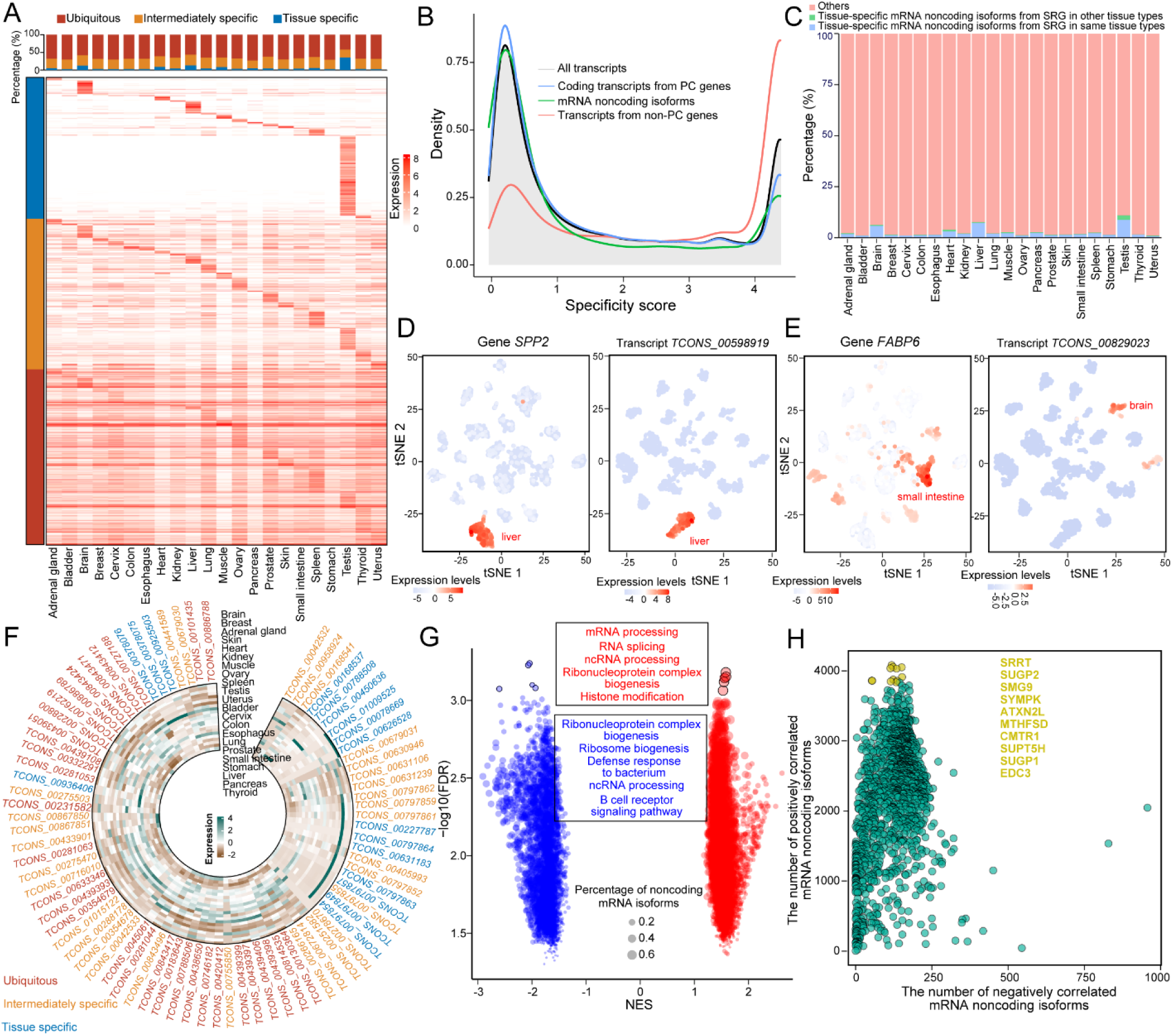
Tissue-specific expression and functional characteristics of mRNA noncoding splice isoforms across human normal tissues. **A.** Heatmap illustrating expression profiles of mRNA noncoding isoforms originated from protein-coding genes across different human normal tissues. **B.** Line plots depicting the density distribution of tissue specificity scores for all transcripts, protein-coding transcripts from protein-coding genes, noncoding transcripts from protein-coding genes, and transcripts from non-protein-coding genes. **C.** Stacked bar plots indicating the proportion of tissue-specific mRNA noncoding isoforms originating from host tissue specific genes (TSGs) across different human normal tissues. **D.** tSNE visualizations of gene *SPP2* and its noncoding isoform *TCONS_00598919* expression levels across different human normal tissues. **E.** tSNE visualizations of gene *FABP6* and its noncoding isoform *TCONS_00829023* expression levels across different human normal tissues. **F.** Circular heatmap showing expression patterns of noncoding isoforms from protein-coding genes associated with embryonic development. **G.** Volcano plot displaying biological processes significantly enriched among protein-coding genes correlated with noncoding isoforms. **H.** Scatter plot presenting the count of positively and negatively correlated noncoding isoforms for each RNA-binding protein (RBP).

### Expression landscape of mRNA noncoding splice isoforms across different human cancer types

According to noncoding isoform expression levels in 9,219 samples across 33 human cancer types and 651 related adjacent normal tissues, we identified 12,156 detectable noncoding isoforms with an average (median) number of TPM > 0 across all tumors and adjacent normal tissues (**Fig. 4A**). The number of noncoding isoforms expressed by each tumor accounts for approximately one-third of the total level. Among these isoforms, we observed that some noncoding isoforms were highly expressed across all cancer types (**Fig. 4B**). Expression similarity analysis revealed a strong cancer-type-specific pattern, as samples from the same cancer type clustered together (**Fig. 4C**). Differentially expressed noncoding splice isoforms (DEI) analysis revealed a higher proportion of upregulated noncoding isoforms in tumors, ranging from 4.48% in UCEC to 30.4% in LIHC (median: 14.6%). Downregulated noncoding isoforms range from 0.37% in ESCA to in 19.9% KICH (median: 6.72%) (**Fig.4D, table S2**). We classified these DEIs into three groups: 561 ubiquitous DEIs in ≥9 cancer types, 8,996 intermediately specific DEIs that are expressed in 2–8 cancer types, and 1,691 cancer-type-specific DEIs that are expressed in only one cancer type (**Fig.4E**). The overall frequency of transcript abundance differences, defined as changes in the proportion of each transcript within the total gene reads, was generally similar to conventional expression level differences based on TPM values (P > 0.05). Notably, seven cancer types, including breast cancer, clear cell renal cell carcinoma, papillary renal cell carcinoma, lung adenocarcinoma, lung squamous cell carcinoma, pancreatic ductal adenocarcinoma, and papillary thyroid carcinoma, exhibited significant transcript abundance variations, indicating complex transcript isoform selection mechanisms. Of differentially expressed noncoding isoforms, an average of 38.8% showed concordant abundance and expression change, while 61.2% displayed asymmetric regulation, with only 2.2% exhibiting an inverse relationship (**Fig. S1**). To assess the clinical relevance of noncoding DEIs, we developed a DEIs-based risk score model for 17 cancers. Gene set enrichment analysis (GSEA) showed that noncoding DEIs correlated positively with cancer hallmarks across multiple cancer types (**Fig. 4F**). Additionally, we divided patients into risk score-low and -high group. Patients with low-risk score demonstrated a prominent survival benefit in multiple cancers (**Fig 4G and Fig S2**). Multivariate Cox regression analysis confirmed that the noncoding DEI risk score was an independent prognostic factor (**Fig 4H**). The noncoding DEI risk scores varied significantly among molecular subtypes, correlated with tumor stages, and were elevated in advanced cancers (**Fig. S3**).

**Fig. 4.**
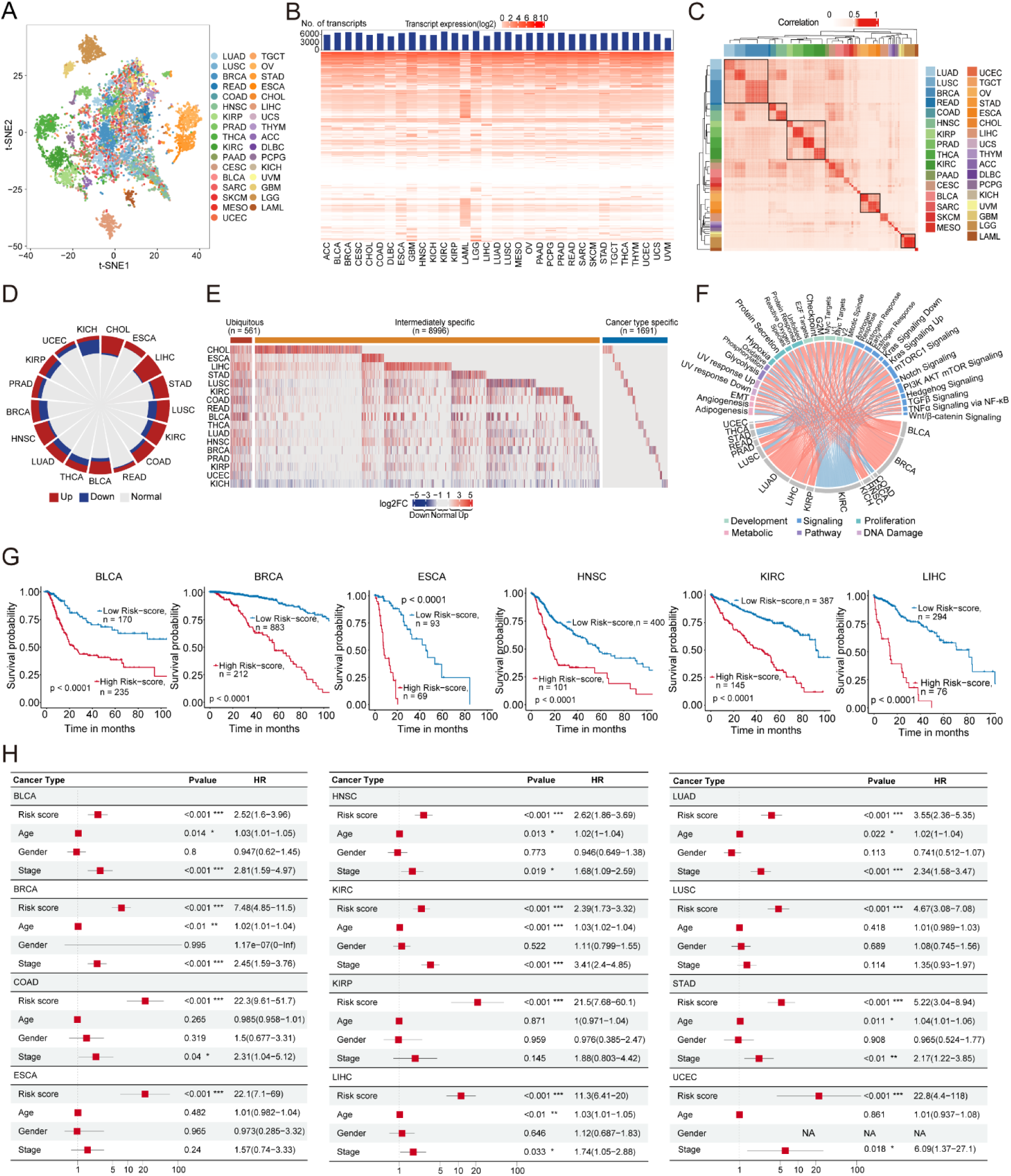
Expression landscape of mRNA noncoding splice isoforms across different human cancer types. **A.** t-SNE projection was used to visualize the expression profiles of 12,156 mRNA noncoding splice isoforms across 33 cancer types. Distinct clustering patterns were observed for several cancer types. **B.** Number of detectable mRNA noncoding splice isoforms across cancer types and their global expression profiles. Each column represents a cancer type, and each row corresponds to an mRNA noncoding splice isoform. Color indicates expression level. **C.** Expression similarity among different tumor samples. The color bar denotes cancer types, and the scale bar indicates the degree of expression similarity among different tumor samples. **D.** Expression changes of mRNA noncoding isoforms (measured by TPM) across different cancers. Red, blue, and gray represent the percentages of upregulated, downregulated, and non-significantly changed isoforms, respectively. **E.** Differentially expression profile of mRNA noncoding splice isoforms among different human cancers. Red, green, and blue bars indicate ubiquitous (≥9), intermediately specific (2–8), and cancer-type-specific DEIs, respectively. **F.** Associations between risk scores and cancer hallmarks. Color bars represent different hallmarks; link thickness corresponds to the Log P-value. Red links indicate positive correlations and blue links indicate negative correlations. **G.** Kaplan–Meier survival curves for patients with high (red) and low (blue) risk scores across different human cancers. **H.** Multivariate Cox regression analysis incorporating risk score, patient age, gender, tumor stage, and clinical outcomes in different human cancers. The center of the red squares represents HR values, and horizontal lines denote the 95% confidence intervals (CI) associated with each variable.

### Polysome profiling and in vitro transcription and translation assays proved noncoding splice isoforms in human mRNA

In the aforementioned studies, we systematically accessed the protein-coding potential of mRNA noncoding splice isoforms using three independent bioinformatics prediction tools. To verify whether these predicted noncoding transcripts possess protein translation capacity, we examined representative candidates from human normal skin, liver, and testicular tissues. Sequence alignment using the Ensembl database displayed the genomic organizations and exon constituents between the candidate noncoding splice isoforms and their corresponding classic protein-coding isoforms (**Fig. S4**). Primers targeting isoform-specific exons were designed for polysome profiling analysis, which showed no detectable polysome associations for the indicated 9 noncoding splice isoforms including EDDM13, C16orf95, TP53AIP1, ACOX2, ERI3, ATP5MPL, NBR1, FOXN3 and TIMD4 (**Fig. 5A-5C**), demonstrating a lack of translation potentials. Furthermore, we performed cell-free in vitro transcription and translation assays. The results verified the absence of detectable protein products or micropeptides from all the indicated 9 noncoding splice isoforms (**Fig. 5D**). Collectively, these findings proved noncoding splice isoforms in human mRNA.

**Fig. 5.**
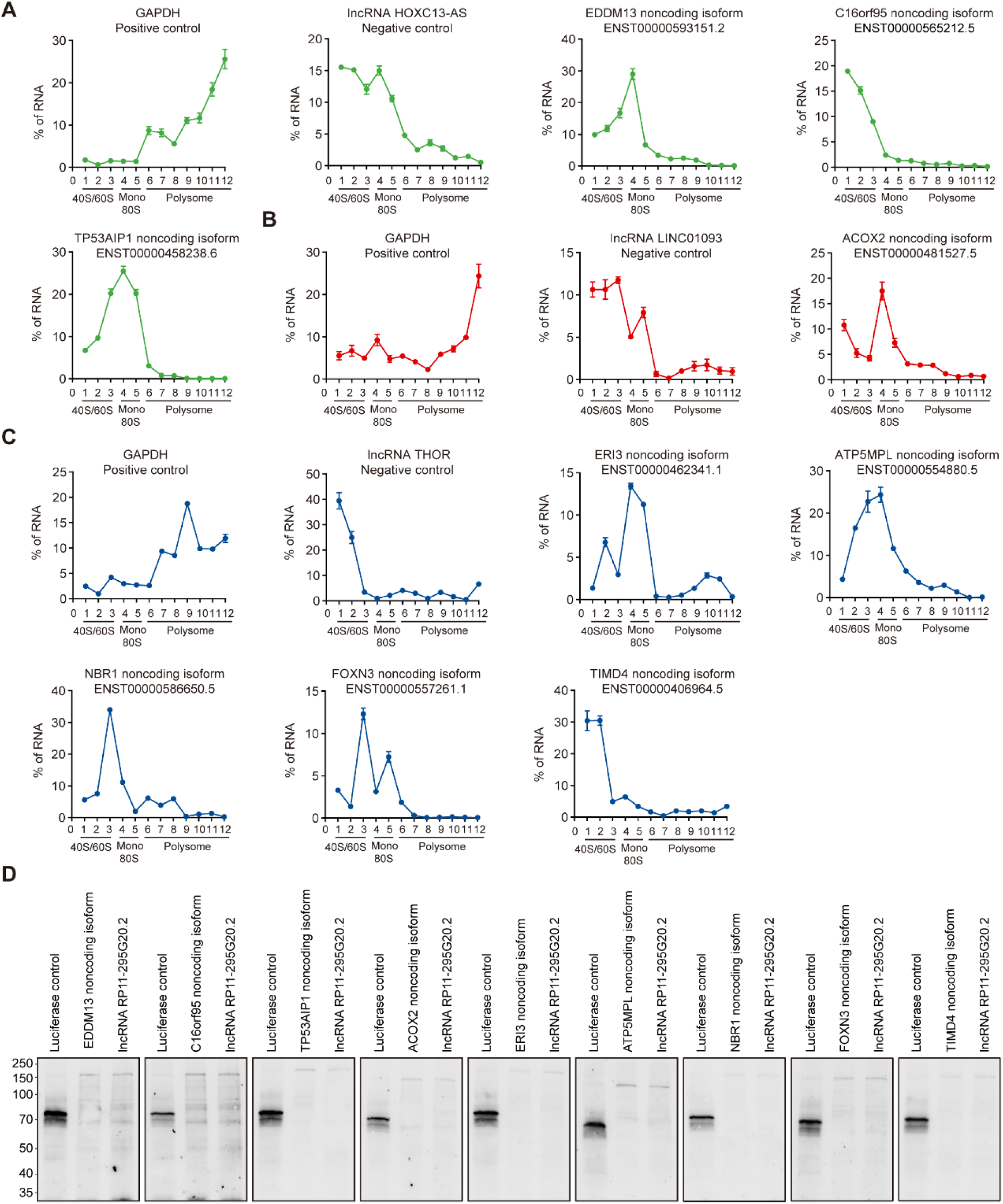
Polysome profiling analyses and in vitro transcription and translation assays proved noncoding splice isoforms in human mRNA. **A.** Polysome profiling analysis of the indicated mRNA noncoding splice isoforms in human skin cell. GAPDH mRNA is used as a positive control for active protein translation; Long noncoding RNA lncRNA HOXC13-AS serves as the negative control. **B.** Polysome profiling analysis of the indicated mRNA noncoding splice isoforms in human liver cell. GAPDH mRNA is used as a positive control for active protein translation; Long noncoding RNA LINC01093 serves as the negative control. **C.** Polysome profiling analysis of the indicated mRNA noncoding splice isoforms in human testicular cell. GAPDH mRNA is used as a positive control for active protein translation; Long noncoding RNA lncRNA THOR serves as the negative control. **D.** In vitro transcription and translation assays of the indicated mRNA noncoding splice isoforms in the cell. Luciferase is used as a positive control for protein synthesis; Long noncoding RNA lncRNA RP11-295G20.2 serves as the negative control.

## Discussion

In this study, we discovered that 7,298 human protein-coding genes produce mRNA noncoding splice isoforms besides protein-coding counterparts in the genome. Alternative splicing and alternative polyadenylation enable dual-output from a single protein-coding gene. The discovery of noncoding splice isoforms adds a new dimension to our understanding of mRNA functional property, and highlights the translation-noncoding duality of mRNA originated from human protein-coding genes. To date, two examples have been reported in other species, i.e. a non-coding CSF-1R isoform regulating germ cell dynamics identified in rats (*20*), and a non-coding IL-4 isoform modulating immune activity identified in mice (*21*). It is unclear whether the translation-noncoding duality of mRNA represents a universal phenomenon across the eukaryotes, particularly in multi-exon organisms, and if so, whether it reflects evolutionarily conserved principles. This study opens a new field for exploring the evolutionary conservation, regulatory mechanisms, and biological significance of mRNA non-coding splice isoforms across diverse organisms.

The generation of mRNA noncoding isoforms represents an intriguing aspect of post-transcriptional regulation due to its potential implications in gene expression and functional diversification. Among identified AS events, alternative first exon (AF) events emerged as the most frequent, suggesting possible novel transcriptional start sites. The utilization of alternative transcriptional start sites could produce distinct 5’ termini, potentially disrupting conventional translation initiation signals or sequences (*22–24*), thereby converting originally protein-coding transcripts into noncoding RNA isoforms. Approximately 30% of identified mRNA noncoding isoforms exhibited high inclusion ratios, indicating frequent incorporation of alternatively spliced exons or introns. This observation raises the possibility that alternative exon or intron usage could disrupt open reading frames (ORFs), potentially leading to frameshift mutations or premature termination codons (*25–27*). Such disruptions would consequently abolish protein-coding potential, resulting in noncoding isoforms. Additionally, noncoding isoforms generally displayed shorter polyadenylated tails relative to other transcripts derived from the same or different protein-coding genes. These shortened poly(A) tails may imply premature polyadenylation, potentially truncating the ORF and hindering translation initiation or completion, thereby producing noncoding isoforms. Notably, certain mRNA noncoding isoforms displayed tissue-specific AS patterns. Tissue-specific factors, including splicing regulators, epigenetic modifiers, transcriptional factors, or RBPs, might preferentially modulate the expression and processing of these isoforms in particular tissue contexts (*28–31*). These regulatory interactions highlight a sophisticated player of gene expression control, underpinning tissue-specific functional specialization and adaptability.

The expression profiles of mRNA noncoding isoforms in normal tissues and during developmental stages exhibit intriguing characteristics that warrant further exploration, emphasizing their potential significance and biological value. Notably, genes positively correlated with these noncoding isoforms were significantly enriched in RNA processing-related pathways, such as “mRNA processing”, “RNA splicing”, and “ncRNA processing”. This enrichment aligns remarkably well with our observation, where most noncoding mRNA isoforms originated from protein-coding genes involved in RNA processing pathways. This alignment suggests potential functional relationships between noncoding isoforms and their protein-coding counterparts, indicating possible interactions or coordinated activities. Such relationships could represent an additional layer of regulatory complexity; wherein noncoding isoforms modulate the functions of their coding isoforms or participate in broader regulatory network to finely tune RNA processing activities. Furthermore, certain tissue-specific noncoding isoforms were derived from tissue-specific genes, either within the same or different tissues, reflecting precise regulatory controls tailored to tissue-specific functions. The distinct tissue-specific expression patterns suggest that these noncoding isoforms may serve specialized roles in maintaining tissue identity or performing unique regulatory functions. Thus, these isoforms could potentially act as biomarkers or “tissue tags”, aiding in tissue-specific diagnostics or therapeutic targeting (*31, 32*). Moreover, the association of mRNA noncoding isoforms with genes involved in human embryonic development suggests their involvement in specialized developmental functions. Their presence underscores the intricately layered regulatory architecture characteristic of eukaryotic development. The developmental-stage-specific noncoding isoforms provided novel regulatory mechanisms essential for precise developmental timing, tissue differentiation, and developmental diversity.

In addition, mRNA noncoding splice isoforms are pervasively presented across human malignancies. An asymmetric regulatory pattern was observed with noncoding splice isoforms more frequently upregulated than downregulated in tumors. While a core subset of noncoding isoforms appears to participate in the common regulatory mechanisms, the prominent cancer-type-specific clustering indicates that some mRNA noncoding isoforms contribute to tissue-specific oncogenic programs. A noncoding DEI-based risk score effectively stratified patient survival across multiple cancer types and acted as an independent prognostic factor. Differentially expressed noncoding splice isoforms were highly correlated with the cancer hallmarks, suggesting the functional involvement in key oncogenic processes during tumorigenesis.

In conclusion, we discovered 15,836 mRNA noncoding splice isoforms across 7,298 protein-coding genes accounting for 36% of total 20, 270 protein-coding genes in human genome. mRNA noncoding splice isoforms display tissue-specific distributions and are generated by alternative splicing and polyadenylation. Most mRNA noncoding isoforms involve in RNA processing pathways. Differentially expressed mRNA noncoding isoforms are associate with the cancer hallmarks and can independently predict patient survival. These findings unraveled human mRNA harbors widespread noncoding splice isoforms, and highlight the translation-noncoding duality of mRNA, providing a new dimension to our understanding of mRNA functional property.

## Materials and Methods

### Data collection

We obtained full-length transcripts expression data for 33 cancer types and 17 matched adjacent normal tissues, and GTEx normal tissues from FLIBase (http://www.flibase.org/) (*16*). Clinical phenotype data were retrieved from Xena (https://xenabrowser.net/datapages/).

### Computational Pipeline for the Selection and Functional Annotation of mRNA noncoding splice isoforms

403,693 full-length transcript isoforms expressed in human normal tissues were obtained from the FLIBase database, based on third-generation long-read sequencing technologies. Transcripts with no detectable expression in normal tissues were excluded. Briefly, transcripts with a median TPM > 0 in at least one normal tissue were retained. Genomic coordinates were assigned using the Ensembl GRCh38 annotation, and only transcripts located within protein-coding gene loci were included. Coding potential was evaluated using three independent tools: CPAT (Coding Potential Assessment Tool) (*17*), CPC2 (Coding Potential Calculator 2) (*18*), and GeneMark (*19*). Transcripts predicted to be non-coding potential by at least two of the three tools were retained for downstream analysis. To eliminate potential misannotations, Bedtools was used to remove non-coding transcripts that overlapped with annotated protein-coding regions in the reference genome. Gene Ontology (GO) enrichment analysis was performed on the filtered non-coding transcript associated protein-coding genes using the clusterProfiler R package (*33*), with statistical significance determined by an adjusted p-value < 0.05.

### Identification of alternative splicing events

Alternative splicing events (ASEs) were identified using SUPPA2 (*34*) (version 2.4), a computational tool designed for quantifying splicing across transcriptomic datasets. The TPM values of transcripts were input into SUPPA2 to generate the percent spliced-in (PSI) values for each ASE. SUPPA2 categorizes ASEs into seven types: exon skipping (SE), mutually exclusive exons (MX), alternative 5’ splice sites (A5SS), alternative 3’ splice sites (A3SS), retained intron (RI), alternative first exon (AFE), and alternative last exon (ALE). For each event type, SUPPA2 identifies potential splicing isoforms by analyzing transcript structures provided by the annotation (GENCODE v44). The tool calculates PSI values by comparing the relative abundance of transcripts that include or exclude specific splicing events. This enables comprehensive profiling of alternative splicing pattens within and across sample groups.

### Alternative polyadenylation (APA) analysis

Alternative polyadenylation (APA) analyses were performed using DarPars2 (*35*) (Dynamic analysis of Alternative PolyAdenylation from RNA-seq, version 2.1), a computational framework specifically designed to detect and quantify APA events directly from RNA sequencing data. RNA-seq reads were aligned to the human genome reference (GRCh38) using STAR aligner (v2.7.10a), followed by transcript assembly to identify RNA expression patterns. The aligned BAM files served as input for DaPars2, which detects APA sites by modeling read coverage differences in the 3’ untranslated regions (3’UTRs) of genes. DaPars2 estimates the proximal-to-distal polyadenylation usage index (PDUI), representing the relative usage between proximal and distal polyA sites for each gene.

### Calculation of tissue-specific scores

Tissue-specific scores of transcripts were calculated using a method that was described in our previous study (*36*). Specifically, transcript specificity scores were computed by subtracting the Shannon entropy of transcript expression from the logarithm of the total number of tissues analyzed. The formula applied was:

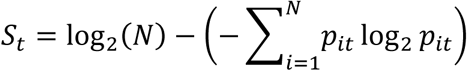

where *S_t_* denotes the specificity score for transcript *t*, *N* is the total number of tissue types, and *p_it_* represents the proportion of transcript *t*’s expression in tissue *i*. Each transcript was thus assigned one specificity score along with *N* expression proportions corresponding to each tissue. These proportions were determined as follows:

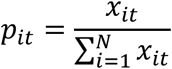

where *x_it_* is the expression level of transcript *t* in tissue *i*. Transcripts were designated tissue-specific if the highest expression proportion was greater than twice the second-highest proportion and if the specificity score exceeded 1.

### Functional enrichment analysis of mRNA noncoding splice isoforms

To identify protein-coding genes correlated with each noncoding splice isoform, Spearman correlation coefficient (*ρ*) were calculated based on their respective expression levels across samples. A correlation score (*S_cor_*) was defined for each pair of noncoding isoform and protein-coding gene using the following formula:

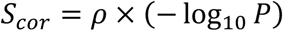

where *S_cor_* represents the correlation score, *ρ* indicates the Spearman correlation coefficient, and *P* denotes the *p*-value obtained from the correlation test. Protein-coding genes were subsequently ranked based on these correlation scores for each noncoding isoform. Gene set enrichment analysis (GSEA) of the ranked lists was performed to identify significantly enriched biological processes, utilizing the clusterProfiler (*37*) R package (version 4.12.6).

### mRNA noncoding splice isoforms filtering in pan-cancer analysis

The initial set of 15,836 noncoding transcript isoforms was further filtered based on expression profiles. Transcript isoforms with no detectable expression in either tumor or adjacent normal tissues were excluded. Briefly, for the 16 cancer types lacking paired normal tissues, transcript isoforms with a median TPM > 0 in at least one cancer type were retained. For the 17 cancer types with matched normal tissues, transcript isoforms with a median expression > 0 in either tumor or normal samples were included. The two resulting transcript isoform sets were merged and duplicates removed, yielding a final set of 12,156 transcript isoforms for subsequent analysis.

### t-distributed stochastic neighbor embedding (t-SNE) analysis

t-SNE analysis was performed using the Rtsne package (version 0.17) in R (version 4.4.0). Prior to analysis, expression data for the 12,156 transcript isoforms were log-transformed to stabilize variance, followed by normalization to a mean of 0 and standard deviation of 1. These preprocessing steps were implemented to enhance data quality and comparability across samples. After preprocessing, t-SNE was applied to project high-dimensional transcript isoform expression data into a low-dimensional space for visualization of sample relationships.

### Differential expression analysis of mRNA noncoding splice isoforms

The “fold change” was calculated as the ratio of median transcript isoform expression between tumor and adjacent normal tissues across the 17 cancer types with paired samples. Differential expression was assessed using the Wilcoxon rank-sum test. Transcript isoforms with a p-value < 0.05 were considered significantly differentially expressed. Among these, transcript isoforms were classified as “upregulated” if the log₂ fold change was > 1, “downregulated” if the log₂ fold change was < –1, and “normal” if the log₂ fold change fell between –1 and 1.

### Transcript isofroms’ abundance analysis

For the transcript isoforms (T1, T2, T3…, Tn) contained in one gene, their TPM values (TPM) were used to calculate the abundance of transcript T1 (Abundance_T1_) as follows:

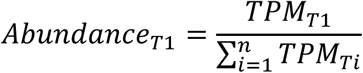

We compared transcript isoform abundance between tumors and adjacent normal tissues, considering a p-value < 0.05 as statistically significant. For each cancer type, transcript isofroms with a mean abundance ratio (tumor to adjacent normal tissues) > 1 were classified as “upregulated,” while those with a ratio < 1 were classified as “downregulated.”

### Calculation of patients’ risk score

To investigate the correlation between differentially expressed transcript isoforms and factors such as tumor hallmarks, cancer subtypes, and patient survival prognosis, we screened 12,156 transcript isoforms (T1, T2, T3…, Tn) with upregulated expression in each of the 17 paired cancer types. We then performed Cox univariate analysis using transcript isoform TPM values (TPM) as the independent variable to calculate the regression coefficient (β). Finally, the risk score was constructed using the following formula:

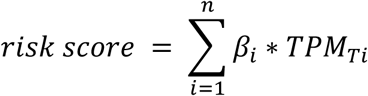

### Gene set variation analysis (GSVA)

We performed GSVA using the R package GSVA (version 1.52.2) to assess gene set variation across hallmark pathways in the 17 paired cancer types. The GSVA score was then used to calculate Spearman’s correlation coefficient with the corresponding patients’ risk scores.

### Survival analysis

We used the R packages survival (version 3.6.4) (R Foundation for Statistical Computing, Vienna, Austria; http://www.r-project.org) and survminer (version 0.4.9) to generate Kaplan-Meier survival curves for 17 paired cancer types. The “surv_cutpoint()” function from survminer was employed to determine the optimal cut-off point for patients’ risk scores. Based on this threshold, patients were stratified into high- and low-risk groups, and survival differences were evaluated using log-rank tests. Univariate and multivariate Cox proportional hazards regression analyses were then performed using the ezcox package (version 1.0.4) to estimate hazard ratios (HRs) and corresponding p-values across cancer types.

### Cell lines and cell culture

Immortalized liver cell line THLE-2 and Human testicular support cells-Immortalized were obtained from the Shanghai Zhong Qiao Xin Zhou Biotechnology Co.,Ltd. NHEK, the human normal epidermal keratinocytes, were kindly provided by Dr. Wei Li (Shanghai Jiao Tong University). THLE-2 cells were cultured in BEGM medium with 10% fetal bovine serum (FBS, Gibco). Human testicular support cells-Immortalized was maintained in Human testicular support cells-Medium for immortalization (Shanghai Zhong Qiao Xin Zhou Biotechnology Co.,Ltd). NHEK cells were cultured in DMEM (Gibco) supplemented with 10% FBS. All cells were grown in 5% CO_2_ at 37°C. All of the cell lines were regularly tested for Mycoplasma free.

### Clone construction, Reverse transcription and Real-time PCR

The full-length cDNA sequences of EDDM13, C16orf95, TP53AIP1, ACOX2, ERI3, ATP5MPL, NBR1, FOXN3 and TIMD4 were individually cloned into the pMD19-T Vector (TaKaRa, Japen), and then subjected to DNA sequencing. All primer sequences used in this study are listed in **Table S3.**

Total RNA from indicated cell lines was isolated with TRIzol (Life Technologies, CA, USA) following the manufacturer’s guidelines. Subsequently, 5 μl RNA was utilized to synthesize complementary DNA (cDNA) deploying PrimeScript^TM^ RT Reagent kit (TaKaRa, Tokyo, Japan). Real-time qPCR was performed using SYBR® Green Pro Taq HS (Accurate Biology, Changsha, China) in 7900 Real-Time PCR apparatus (Applied Biosystems, USA). GAPDH was used as internal control for normalization. Primers used for RT-qPCR in this study were listed in the **Table S3**.

### Polysome profiling analysis

Indicated cells were pretreated with 100 μg/ml cycloheximide (CHX) at 37°C for 10 min, and then 3×10^7^ cells were lysed in lysis buffer (10 mM Tris-HCl pH7.5, 10 mM MgCl_2_, 10 mM NaCl, 1% TritonX-100, 1 mM DTT, 100 μg/ml CHX, 1% sodium deoxycholate, 40U/ml RNase Inhibitor) for 10 min on the ice and then centrifuged at 12,000 g at 4°C for 10 minutes. 800 μl of extract was loaded on 15% to 50% sucrose gradients followed by ultracentrifugation with a SW41rotor (Beckman) at 274,000 g at 4°C for 2 hours. Absorbance at 254 nm was recorded. The corresponding isolated fractions were further used for RNA extraction. The extracted RNA was then used for RT-qPCR analysis.

### In vitro transcription and translation assay

In vitro transcription and translation assay of indicated transcripts were performed deploying a TnT Quick Coupled Transcription/Translation Kit (Promega, USA) and detection was conducted using a Transcend Non-Radioactive Translation Detection System (Promega, USA) according to the manufacturer’s instructions. Briefly, a 50 μl translation reactions including TnT T7 Quick Master Mix, 1mM Methionine, Plasmid DNA Template (0.5 μg), Transcend^TM^ Biotin-Lysyl-tRNA, T7 TnT^®^ PCR Enhance, and Nuclease-Free Water was incubated at 30°C for 90 minutes. The translated proteins were then detected by HRP Streptavidin (YEASEN, China), which binds biotinylated lysine incorporated into the translated proteins. The luciferase T7 control DNA provided by manufacturer was utilized as a positive control.

### Statistical analysis

Statistical analysis and data visualization in this study were performed by using R software. Unless otherwise specified, all tests were two-tailed, and a P or FDR value < 0.05 was considered to indicate statistical significance.

## Supporting information

Supporting Information including 4 supplemental figures

