## Supporting Information including 4 supplemental figures for "Human Messenger RNA Harbors Widespread Noncoding Splice Isoforms"

#### Figure S1

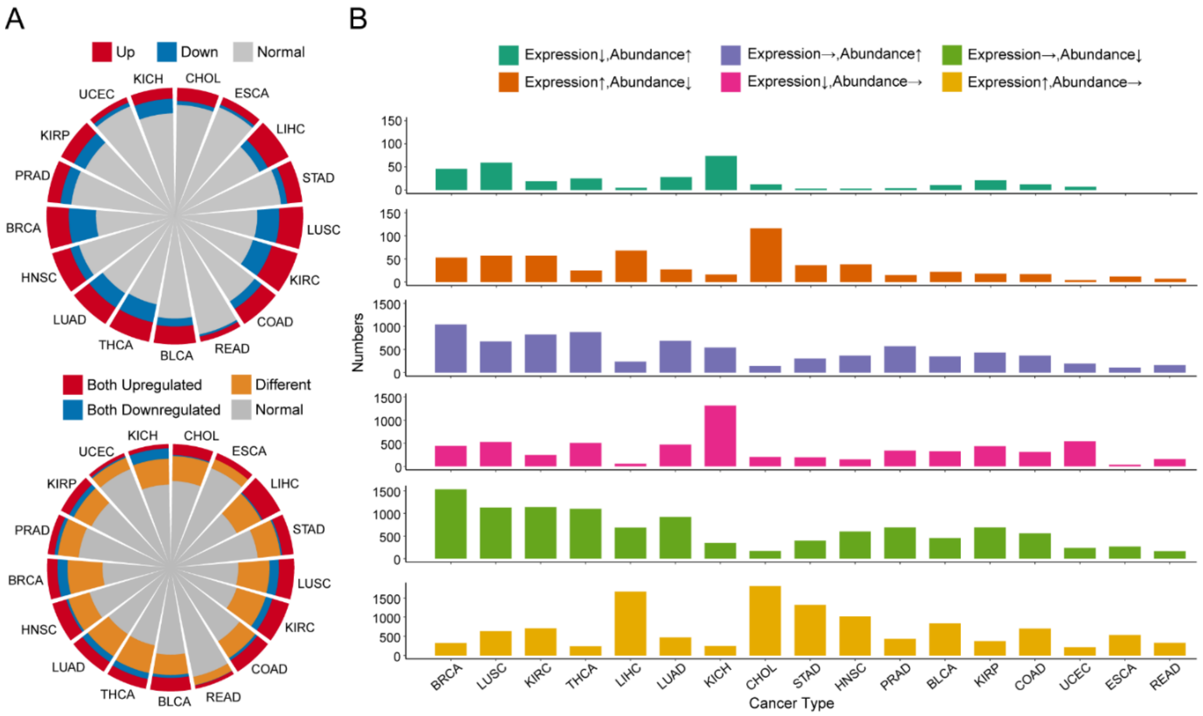


**Figure S1. Expression landscape of mRNA noncoding splice isoforms in human cancer**

**A**. Top: Expression changes of mRNA noncoding splice isoforms based on their abundance across different cancer types. Red, blue, and gray indicate the percentage of upregulated, downregulated, and non-significantly changed noncoding splice isoforms, respectively. Bottom: Overlap of differentially expressed noncoding splice isoforms (DEIs) based on TPM and abundance. Red, blue, orange, and gray represent noncoding splice isoforms that are both upregulated, both downregulated, exhibit asymmetric regulation patterns, or show no significant changes, respectively. **B**. Number of noncoding splice isoforms showing asymmetric regulation patterns in each cancer type. CHOL: Cholangiocarcinoma, ESCA: Esophageal carcinoma, LIHC: Liver hepatocellular carcinoma, STAD: Stomach adenocarcinoma, LUSC: Lung squamous cell carcinoma, KIRC: Kidney renal clear cell carcinoma, COAD: Colon adenocarcinoma, READ: Rectum adenocarcinoma, BLCA: Bladder Urothelial Carcinoma, THCA: Thyroid carcinoma, LUAD: Lung adenocarcinoma, HNSC: Head and Neck squamous cell carcinoma, BRCA: Breast invasive carcinoma, PRAD: Prostate adenocarcinoma, KIRP: Kidney renal papillary cell carcinoma, UCEC: Uterine Corpus Endometrial Carcinoma, KICH: Kidney Chromophobe.

### Figure S2

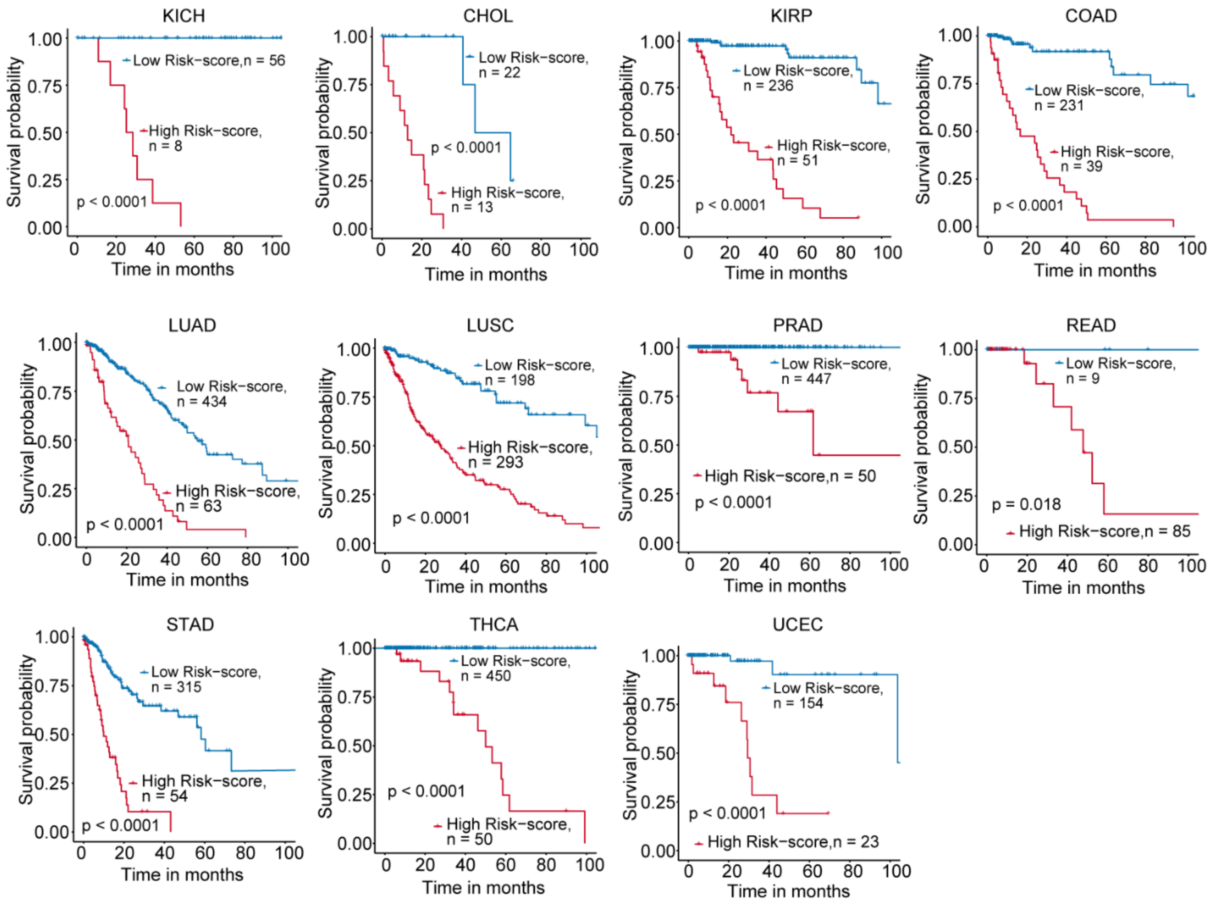


**Figure S2. Noncoding DEIs-based risk score associated with survival in cancer patients.**

Kaplan-Meier survival curves for patients with high (red)and low (blue) risk scores across cancers. The surv_cutpoint() function in the survminer package was used to identify the optimal cut-off point for patient’s risk scores. Patients were then stratified based on this cutoff point, and log-rank tests were conducted to assess differences between groups. KICH: Kidney Chromophobe, CHOL: Cholangiocarcinoma, KIRP: Kidney renal papillary cell carcinoma, COAD: Colon adenocarcinoma, LUAD: Lung adenocarcinoma, LUSC: Lung squamous cell carcinoma, PRAD: Prostate adenocarcinoma, READ: Rectum adenocarcinoma, STAD: Stomach adenocarcinoma, THCA: Thyroid carcinoma, UCEC: Uterine Corpus Endometrial Carcinoma

**Figure S3**

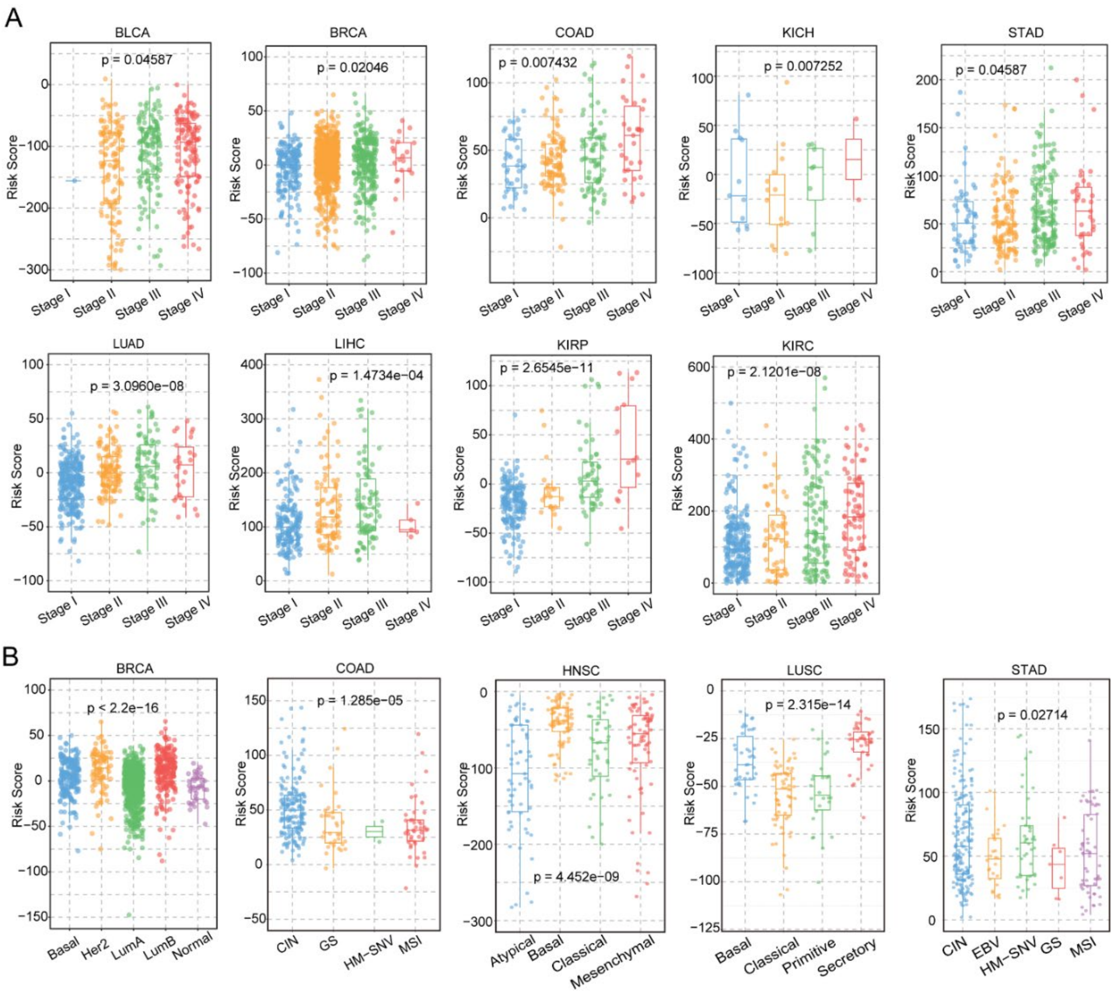


**Figure S3. Noncoding DEIs-based risk score correlated with cancer subtypes and stages.**

A. Association between patient risk scores and cancer stages across 17 cancer types. P-values were calculated using the Kruskal-Wallis H test; only cancer types with $p<0.05$ are shown. B. Association between patient risk scores and molecular or histological subtypes of the 17 cancer types. The points represent each patient's risk score. The box plots show the distribution of all patients' risk scores within each stage or subtype. P-values were also determined by the Kruskal-Wallis H test, and only significant results ($p<0.05$) are presented. BLCA: Bladder Urothelial Carcinoma, BRCA: Breast invasive carcinoma, COAD: Colon adenocarcinoma, HNSC: Head and Neck squamous cell carcinoma, KICH: Kidney Chromophobe, KIRC: Kidney renal clear cell carcinoma,

KIRP: Kidney renal papillary cell carcinoma, LIHC: Liver hepatocellular carcinoma, LUAD: Lung adenocarcinoma, LUSC: Lung squamous cell carcinoma, STAD: Stomach adenocarcinoma.

#### Figure S4

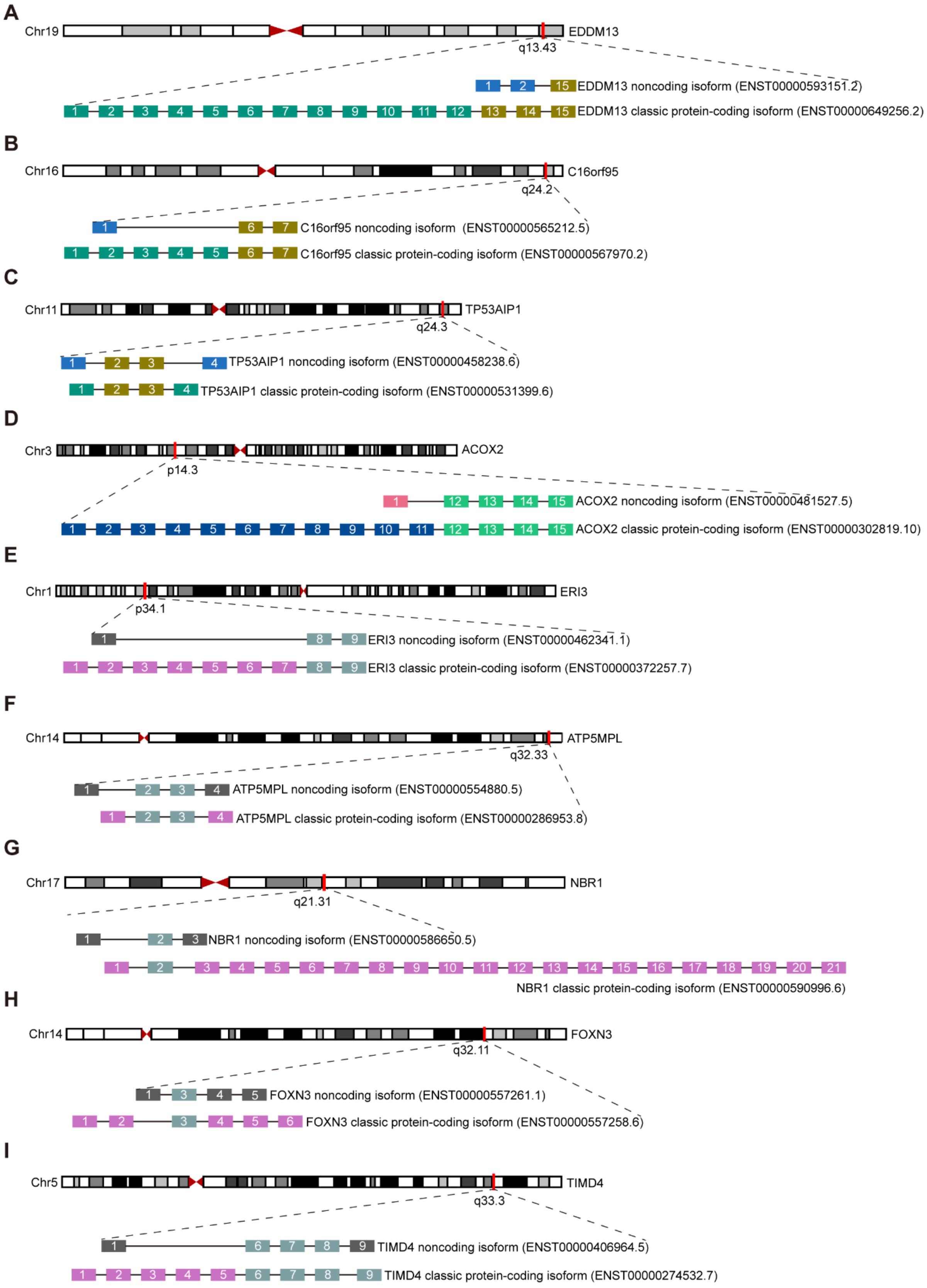
A
Chr19
EDDM13
q13.43
EDDM13 noncoding isoform (ENST00000593151.2)
EDDM13 classic protein-coding isoform (ENST00000649256.2)
B
Chr16
C16orf95
q24.2
C16orf95 noncoding isoform (ENST00000565212.5)
C16orf95 classic protein-coding isoform (ENST00000567970.2)
C
Chr11
TP53AIP1
q24.3
TP53AIP1 noncoding isoform (ENST00000458238.6)
TP53AIP1 classic protein-coding isoform (ENST00000531399.6)
D
Chr3
ACOX2
p14.3
ACOX2 noncoding isoform (ENST00000481527.5)
ACOX2 classic protein-coding isoform (ENST00000302819.10)
E
Chr1
ERI3
p34.1
ERI3 noncoding isoform (ENST00000462341.1)
ERI3 classic protein-coding isoform (ENST00000372257.7)
F
Chr14
ATP5MPL
q32.33
ATP5MPL noncoding isoform (ENST00000554880.5)
ATP5MPL classic protein-coding isoform (ENST00000286953.8)
G
Chr17
NBR1
q21.31
NBR1 noncoding isoform (ENST00000586650.5)
NBR1 classic protein-coding isoform (ENST00000590996.6)
H
Chr14
FOXN3
q32.11
FOXN3 noncoding isoform (ENST00000557261.1)
FOXN3 classic protein-coding isoform (ENST00000557258.6)
I
Chr5
TIMD4
q33.3
TIMD4 noncoding isoform (ENST00000406964.5)
TIMD4 classic protein-coding isoform (ENST00000274532.7)

**Figure S4. Schematic diagrams show the genomic organizations and exon constituents of noncoding splice isoforms and their corresponding classic protein-coding isoforms.**

**A.** Schematic diagram shows the genomic organizations of EDDM13 noncoding splice isoform (ENST00000593151.2) and its corresponding classic protein-coding isoform (ENST00000649256.2) on chromosome Chr19q13.43; **B**. Schematic diagram shows the genomic organizations of C16orf95 noncoding splice isoform (ENST00000565212.5) and its corresponding classic protein-coding isoform (ENST00000567970.2) on chromosome Chr16q24.2; **C**. Schematic diagram shows the genomic organizations of TP53AIP1 noncoding splice isoform (ENST00000458238.6) and its corresponding classic protein-coding isoform (ENST00000531399.6) on chromosome Chr11q24.3; **D**. Schematic diagram shows the genomic organizations of ACOX2 noncoding splice isoform (ENST00000481527.5) and its corresponding classic protein-coding isoform (ENST00000302819.10) on chromosome Chr3p14.3; **E**. Schematic diagram shows the genomic organizations of ERI3 noncoding splice isoform (ENST00000462341.1) and its corresponding classic protein-coding isoform (ENST00000372257.7) on chromosome Chr1p34.1; **F**. Schematic diagram shows the genomic organizations of ATP5MPL noncoding splice isoform (ENST00000554880.5) and its corresponding classic protein-coding isoform (ENST00000286953.8) on chromosome Chr14q32.33; **G**. Schematic diagram shows the genomic organizations of NBR1 noncoding splice isoform (ENST00000586650.5) and its corresponding classic protein-coding isoform (ENST00000590996.6) on chromosome Chr17q21.31; **H**. Schematic diagram shows the genomic organizations of FOXN3 noncoding splice isoform (ENST00000557261.1) and its corresponding classic protein-coding isoform (ENST00000557258.6) on chromosome Chr14q32.11; **I**. Schematic diagram shows the genomic organizations of TIMD4 noncoding splice isoform (ENST00000406964.5) and its corresponding classic protein-coding isoform (ENST00000274532.7) on chromosome Chr5q33.3.
